# GDAP1 binds 4-hydroxynonenal, the toxic end-product of lipid peroxidation, using its GST-like binding pocket

**DOI:** 10.1101/2022.10.28.514248

**Authors:** Matthew R. Googins, Maya Brown, Aigbirhemwen O Woghiren-Afegbua, Kirill I. Kiselyov, Andrew P. VanDemark

## Abstract

GDAP1 (Ganglioside-induced differentiation-associated protein 1) is a novel member of the GST superfamily of detoxifying enzymes that is anchored to the outer mitochondrial membrane. GDAP1 mutations and changes in expression levels result in the inherited neuropathy Charcot-Marie-Tooth (CMT) disease, types 2K, 4A and 4H. GDAP1 activity has been associated with many mitochondrial functions however direct molecular interactions underpinning these connections have remained elusive. Here we establish that GDAP1 can bind 4-hydroxynonenal (4HNE), a toxic end-product of lipid peroxidation. 4HNE binding requires the α-loop, a large sequence motif that is inserted within the substrate recognition domain and is unique to GDAP1. In human cells, GDAP1 overexpression plays a cytoprotective role against oxidative stress. This effect is lost upon deletion of the α-loop. Lastly, we demonstrate that a CMT-causing mutant that destabilizes α-loop positioning also results in a decrease in 4HNE binding affinity. Together these results establish 4HNE as the biological ligand for GDAP1, provide mechanistic insight into 4HNE binding, and demonstrate that altered 4HNE recognition is the likely mechanism underlying CMT-causing mutants such as T157P near the 4HNE binding site.

## Introduction

Oxidative stress is a condition caused by the imbalance between the production of reactive oxygen species (ROS) and the cell’s capacity to neutralize reactive intermediates and repair the molecular damage they generate (Pizzino et al., 2017). Interactions between ROS, especially hydroxyl and peroxide radicals, and polyunsaturated fatty acids can produce fatty acid radicals which initiate a cascade of reactions known as lipid peroxidation that eventually result in the formation of the reactive aldehydes malondialdehyde (MDA) and 4-hydroxy-2-nonenal (4HNE) (Esterbauer et al., 1991). These reactive end products of lipid peroxidation are cytotoxic: MDA can react with guanosine bases in DNA to form the mutagenic DNA adduct M1dG (Marnett, 1999), while both MDA and 4HNE can form Michael adducts or Schiff bases with thiol and amine groups within proteins (Ayala et al., 2014; Esterbauer et al., 1991). 4HNE adduct formation has been shown to inactivate many proteins including cytochrome c oxidase and reductase (Chen et al., 1998; Hwang et al., 2020), and treatment of cells with 4HNE has demonstrated that hundreds of cellular proteins are sensitive to 4HNE adduct formation (Roe et al., 2007; Vila et al., 2008).

Ganglioside-induced differentiation-associated protein 1 (GDAP1) is an emerging member of the glutathione-S-transferase (GST) superfamily of cytoprotective enzymes. GDAP1 is localized to the outer mitochondrial membrane via a C-terminal membrane anchoring sequence (Wagner et al., 2009), and is highly expressed in peripheral neurons (Niemann et al., 2005; Noack et al., 2012; Pedrola et al., 2008; Pedrola et al., 2005), and mutants in *GDAP1* result in types 2K, 4A and 4H of Charcot-Marie-Tooth (CMT), the most common inherited peripheral neuropathy (Barreto et al., 2016; Theadom et al., 2019). GDAP1 has been proposed to play several roles in mitochondrial physiology including mitochondrial network dynamics and transport (Cantarero et al., 2021; Civera-Tregon et al., 2021; Niemann et al., 2005; Niemann et al., 2009), calcium homeostasis (Gonzalez-Sanchez et al., 2019), and oxidative stress (Miressi et al., 2021; Niemann et al., 2014; Noack et al., 2012). GDAP1 knockdown favors mitochondrial elongation (Niemann et al., 2005) and increases the sensitivity of cells to oxidative stress, while overexpression is associated with mitochondrial fragmentation (Niemann et al., 2005; Niemann et al., 2009) and increased glutathione levels (Noack et al., 2012). These results suggest that the function of GDAP1 impacts mitochondrial dynamics and cellular redox state but a molecular mechanism connecting GDAP1 with these functions has remained elusive.

Canonical GSTs conjugate glutathione (GSH) with the electrophilic center of hydrophobic co-substrates (Oakley, 2011; Reinemer et al., 1991) including xenobiotics and oxidized lipids, facilitating their transport out of the cell (Hauck and Bernlohr, 2016; Hayes et al., 2005; Hayes et al., 1998; Hayes and Strange, 1995). Primary sequence analysis indicates GDAP1 contains two GST domains: a putative glutathione (GSH) binding domain (called the G-site), and an H-site responsible for recognizing hydrophobic substrates in canonical GSTs (Awasthi et al., 1993; Hayes et al., 2005). Previous structural examination of the GST-like core of GDAP1 demonstrated that while the G-site still adopts a thioredoxin fold, alterations in putative GSH-interacting residues suggest GDAP1 cannot recognize GSH in the canonical orientation (Googins et al., 2020). This is supported by numerous biochemical observations (Googins et al., 2020; Huber et al., 2016; Shield et al., 2006; Sutinen et al., 2022) using purified components. Observations of GDAP1 H-site structure have shown that it adopts a fold consistent with the GST family (Googins et al., 2020; Nguyen et al., 2020; Sutinen et al., 2022), suggesting that it may be capable of binding GST substrates. Screening for biological small molecules identified thapsic acid (also known as hexadecanedioic acid) as a lipid binding partner (Nguyen et al., 2020). Interestingly, structural characterization of the GDAP1-thapsic acid complex suggests this interaction does not utilize the canonical GST binding pocket, but instead is housed within a separate pocket in the H-site formed by loops near helices 5 and 7 (Nguyen et al., 2020). Thapsic acid is found within the outer mitochondrial membrane (Pettersen and Aas, 1974) but has not been reported to be a GST substrate, further supporting the hypothesis that GDAP1 functions as a lipid sensitive sensor or receptor (Googins et al., 2020; Nguyen et al., 2020). Alternatively, thapsic acid binding may be independent of other binding activities that are housed within the canonical GST binding pocket.

We have demonstrated that GDAP1 can bind the pan-GST inhibitor ethacrynic acid (EA) which is known to target the canonical GST binding pocket through both biochemical and structural studies (Awasthi et al., 1996; Awasthi et al., 1993; Cameron et al., 1995; Googins et al., 2020). Interestingly, EA binding required the α-loop region of GDAP1 (amino acids 145-200), an insertion within the H-site that is unique to GDAP1 (Estela et al., 2011; Googins et al., 2020; Marco et al., 2004; Shield et al., 2006). Existing data indicate that 1) the α-loop is all or partially disordered structurally, indicating that there is inherent flexibility in this region of the protein (Googins et al., 2020; Nguyen et al., 2020); 2) one orientation of the α-loop extends long helices 4 and 5 in a conformation similar to the “tower” domain of lignin (Helmich et al., 2016; Nguyen et al., 2020); 3) prediction of GDAP1 structure from AlphaFold suggest the α-loop may adopt a two helical bundle with hinges that allow it to fold over the G- and H-sites, making a direct contact with the G-site and forming a lid that completes the canonical binding pocket (Jumper and Hassabis, 2022), and 4) the α-loop domain contains a number of CMT causing mutations that are found either within the helical bundle but also near the hinge regions. Thus while GDAP1 appears similar to other GST enzymes in many respects, the identity of a GST-like ligand/substrate and the role of the α-loop in GDAP1 function are both unknown.

Here we show that GDAP1 specifically recognizes 4HNE, a reactive lipid aldehyde that, in addition to a pathological role in oxidative stress, is now recognized as a lipid second messenger (Bae et al., 2011; Zarkovic et al., 1999; Zhang and Forman, 2017). This suggests a direct role for GDAP1 in recognizing the products of oxidative stress. We reveal that in addition to binding, GDAP1 can form covalent adducts with 4HNE which can be distinguished from non-covalent binding events. We show that like ethacrynic acid, 4HNE binding requires the α-loop. We find that binding of thapsic acid cannot compete with 4HNE binding suggesting these activities are very likely housed at different locations on the GDAP1 surface. Using x-ray crystallography, we report the structure of GDAP1 containing a T157P mutation found in patients containing CMT4K. The mutant appears to destabilize the α-loop, perhaps by decoupling motions of the lids from binding events canonical GST pocket. In support of this, we observe a >3-fold reduction in 4HNE binding affinity using GDAP1T157P. Together these findings establish 4HNE as a biologically relevant interaction partner for the canonical GST binding pocket that connects GDAP1 function to products of oxidative stress.

## Results

### GDAP1 regulates cellular redox status

GDAP1 was previously implicated in the regulation of cellular redox (Niemann et al., 2014). We confirmed that GDAP1 has an impact on cellular redox status and that GDAP1 overexpression protects the cells against oxidative stress, using two assays: AlamarBlue and WST-1. AlamarBlue contains resazurin, a non-florescent cell-permeable compound that becomes fluorescent when it is reduced to resorufin (O’Brien et al., 2000). A resorufin buildup in the presence of resazurin is an indicator of reducing activity in the cytoplasm. **Figure 1A** shows that HEK293 cells exposed to AlamarBlue accumulate red resorufin fluorescence in a time-dependent manner, and that the rate of this accumulation is suppressed by the exposure of cells to the prooxidant tetra-Butyl Hydroperoxide (tBHP). This suppression is significantly less pronounced in GDAP1 overexpressing cells, compared with control cells (the graph represents 1 trial, 8 biological replicates). **Figure 1B** (4 trials, 8-16 biological replicates each trial) summarizes the effects of GDAP1 overexpression on the ability of cells to withstand redox shifts induced by tBHP. In control cells, a range of tBHP concentrations causes a significant loss of AlamarBlue response; but this loss is diminished in GDAP1 overexpressing cells (**Figure 1B**).

**Figure 1.**
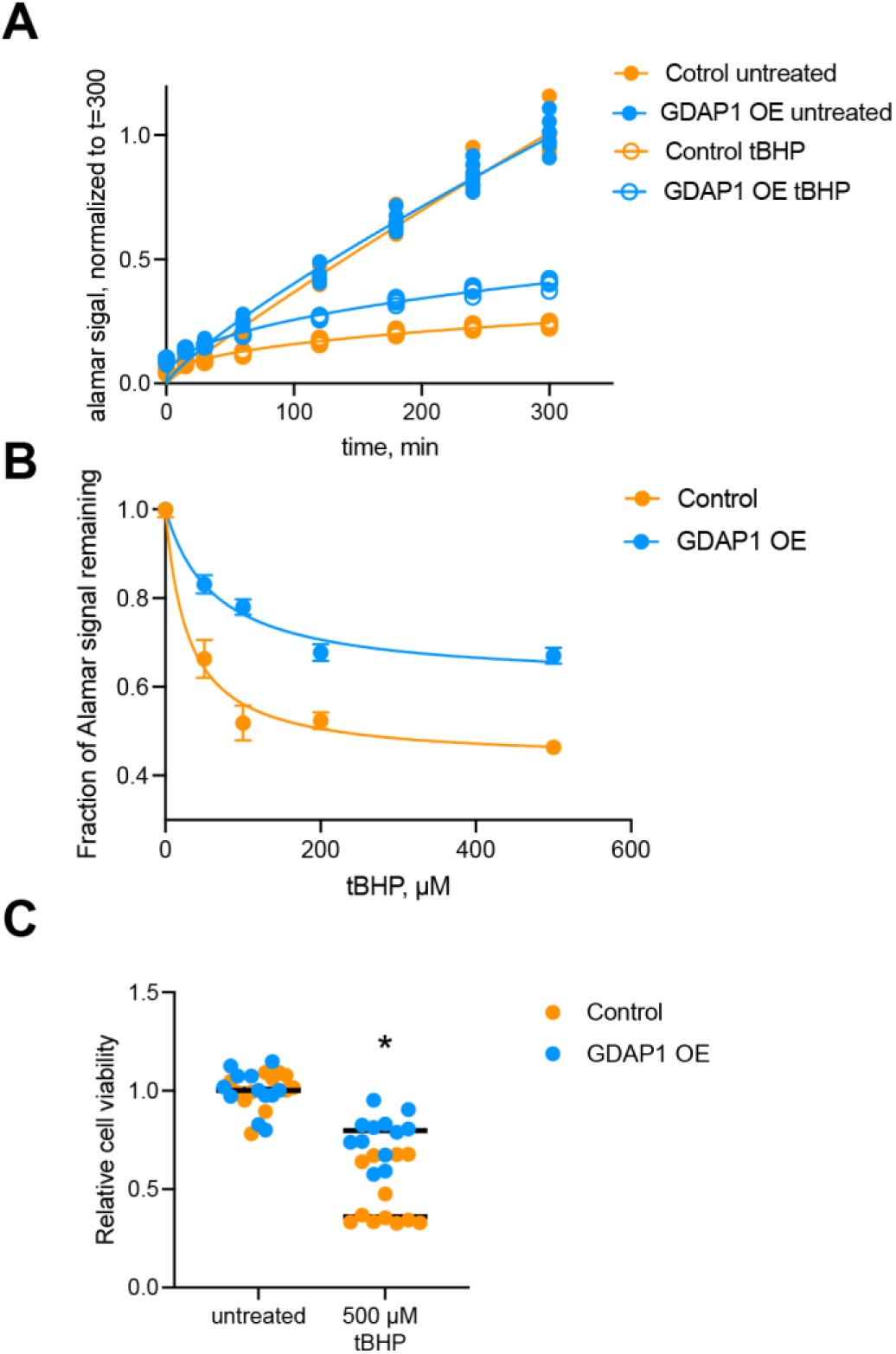
GDAP1 effects cellular responses to the prooxidant tBHP. A) Time course of AlamarBlue signal in HEK293 cells: control and stably transfected with human GDAP1. B) Concentration dependence of tBHP effect on AlamarBlue fluorescence in control and GDAP1 overexpressing cells. In A and B, points represent biological replicates obtained in three independent trials and individual curves demonstrate that the treatment groups are significantly different from each other (see Methods). C) GDAP1 improves cell viability when challenged with 500μM tBHP, as measured using WST-1 assay. Two trials with a total of 12 biological replicates each. * represents p<0.05 by a multiple t test.

WST1 is used to track cell viability, as a measure of mitochondrial dehydrogenase activity (Stockert et al., 2018). **Figure 1C** shows that tBHP suppresses WST-1 signal, which is consistent with decreased cell mitochondrial activity and cell viability. As with AlamarBlue, the suppression is significantly less pronounced in GDAP1-overexpressing cells. Based on these data, we conclude, in line with the prior evidence (Miressi et al., 2021; Niemann et al., 2014; Noack et al., 2012), that GDAP1 has a role protecting cells against oxidative stress.

### GDAP1 binds 4-hydrononenal, the end-product of lipid peroxidation

GDAP1-mediated changes in cellular redox state as well as its primary sequence homology to the GST superfamily of protein suggest that GDAP1 may be capable of binding products of lipid peroxidation. To test this hypothesis, we purified recombinant mouse GDAP1ΔTM (amino acids 1-322) as we have previously (Googins et al., 2020). We then used native PAGE to determine if GDAP1ΔTM could interact with products of lipid peroxidation, focusing on 4HNE and MDA, the two end-products formed by the decomposition of peroxidated lipids (Esterbauer et al., 1991). The migration of GDAP1 does not change in the presence of MDA, suggesting it does not bind, however, we observe the appearance of slower migrating species upon the addition of 1 mM 4HNE (**Figure 2A**), indicating formation of GDAP1-4HNE complexes. The addition of thapsic acid results in a species that migrates slightly faster than GDAP1 alone consistent with previous observations of thapsic acid treatment which was found to reduce GDAP1’s radius of gyration (Sutinen et al., 2022). We also tested arachidonic and linoleic acids to ask whether GDAP1 could recognize fatty acid precursors of lipid peroxidation. We found no observable binding with these ligands, suggesting that the interaction requires the electrophilic groups at the center of 4HNE. Next, we tested binding to trans-2-nonenal (t-2NE) which is similar to 4HNE but does not contain a hydroxyl at the C4 position (**Figure 2B**). Interestingly, we find a reduction in the amount of the bound species with GDAP1 in the presence of t-2NE, (**Figure 2A**), suggesting that GDAP1 is making a direct interaction with the C4 hydroxyl of 4HNE. In an effort to quantify the binding affinity, we measured complex formation throughout a 4HNE titration series, observing an apparent K_D_ of 398 +/− 80 μM (**Figure 2C and D**). The membrane concentrations of 4HNE have been observed to reach as high as 4 mM in the under conditions of oxidative stress (Esterbauer et al., 1991; Koster et al., 1986; Poli et al., 2008) leading us to conclude that GDAP1 is recognizing 4HNE at a biologically relevant concentration. Together, these data suggest that GDAP1 can specifically recognize 4HNE, establishing a new biochemical activity for GDAP1 that directly connects GDAP1 with a known biomarker of oxidative stress (Zarkovic, 2003).

**Figure 2.**
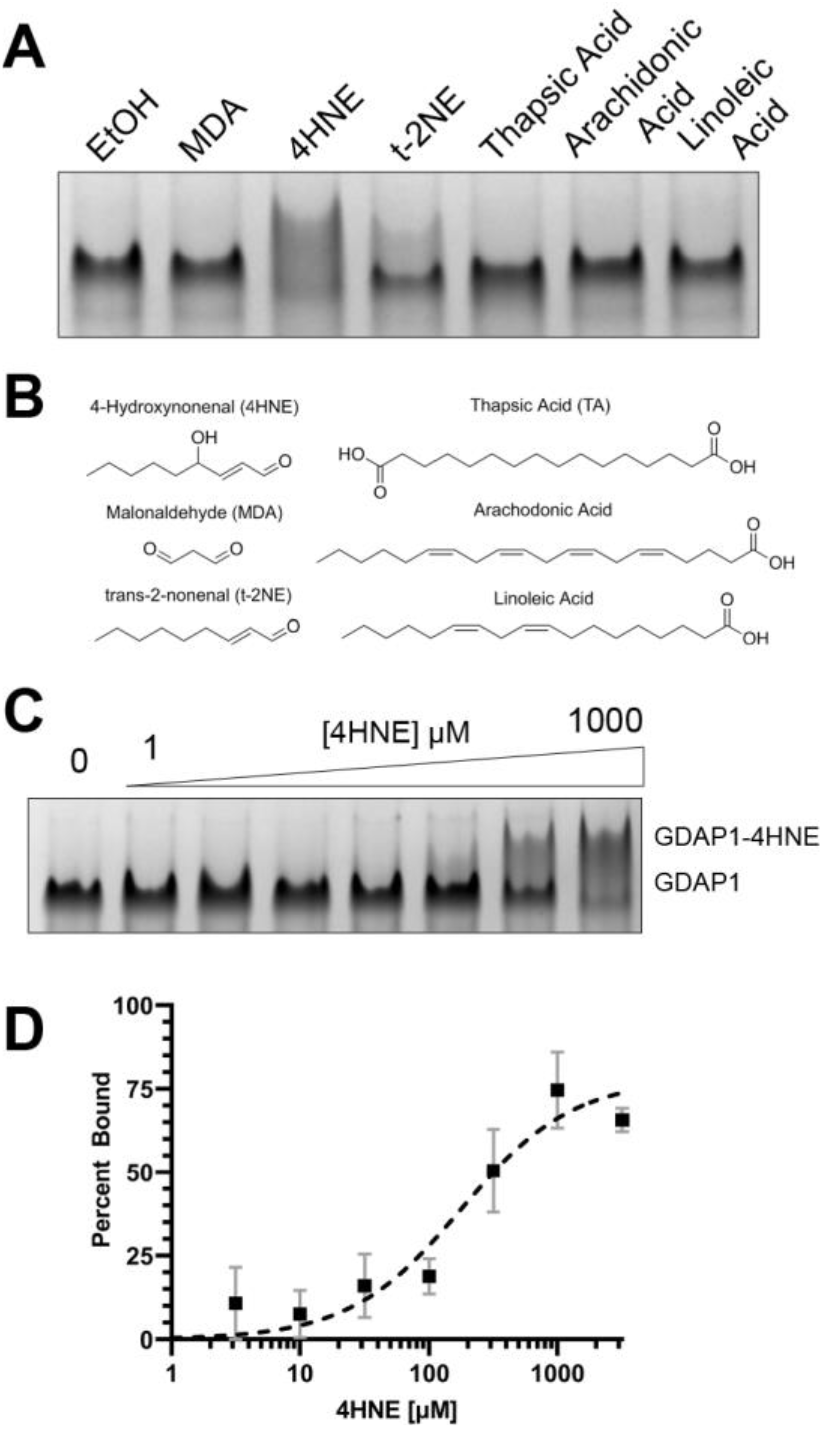
GDAP1 binds 4-hydroxynoneal. **A)** Binding reactions with GDAP1 and indicated fatty acids or products of oxidative stress. Reactions run on 12% Native PAGE. **B)** Chemical structures of compounds used in this study. **C)** Representative gel of GDAP1 binding reactions titrating 4HNE. **D)** Quantification of 4HNE titrations. Error bars represent standard deviation from 3 independent experiments.

### Binding to 4HNE is distinct from 4HNE adduct formation

Increased levels of 4HNE resulting from oxidative stress can result in 4HNE adduct formation, a mechanism known to damage and inactivate proteins, especially those involved in energy production within the mitochondria (Hwang et al., 2020). We first asked whether exposure to 4HNE can produce similar covalent adducts with GDAP1 using GDAP1ΔTM. Reactions containing either GDAP1ΔTM or cofilin1 were incubated with 1 mM 4HNE as in Figure 2, but probed via Western blot using an antibody that specifically recognizes the 4HNE adduct formed by Michael addition to a protein (Usatyuk et al., 2006). Cofilin1 was tested as this protein was proposed to be a GDAP1 binding partner whose binding to GDAP1 was influenced by the cellular redox state (Wolf et al., 2022). We do not observe any 4HNE adduct formation for Cofilin-1, however, we do observe robust 4HNE adduct formation for GDAP1ΔTM (**Figure 3A**). To determine the EC_50_ of this interaction, we monitored adduct formation on GDAP1 as a function of 4HNE titration (**Figure 3C and D**). We find that the EC_50_ for this reaction is 241 ± 52 μM. Since the concentration of 4HNE needed for half maximal adduct formation is similar to that needed for the binding events observed via PAGE, we asked whether the slower migrating species observed via native PAGE represented a covalent 4HNE protein adduct or alternatively a traditional non-covalent protein-ligand interaction. To test this, we performed a 4HNE binding reaction and monitored the results using both native PAGE and Western blot over the course of 1 hour. As shown in **Figure 3B**, binding of 4HNE is observed within the first time point and the relative concentrations of bound and unbound species were unchanged, as would be expected for a binding reaction that has reached equilibrium. These same time points analyzed by Western blot showed that while robust adduct formation was possible, formation of GDAP1-4HNE adducts was not immediate but rather lagged far behind binding. Therefore we conclude that we can observe two distinct species: a 4HNE adduct which is preceded in time by a canonical protein-ligand complex.

**Figure 3.**
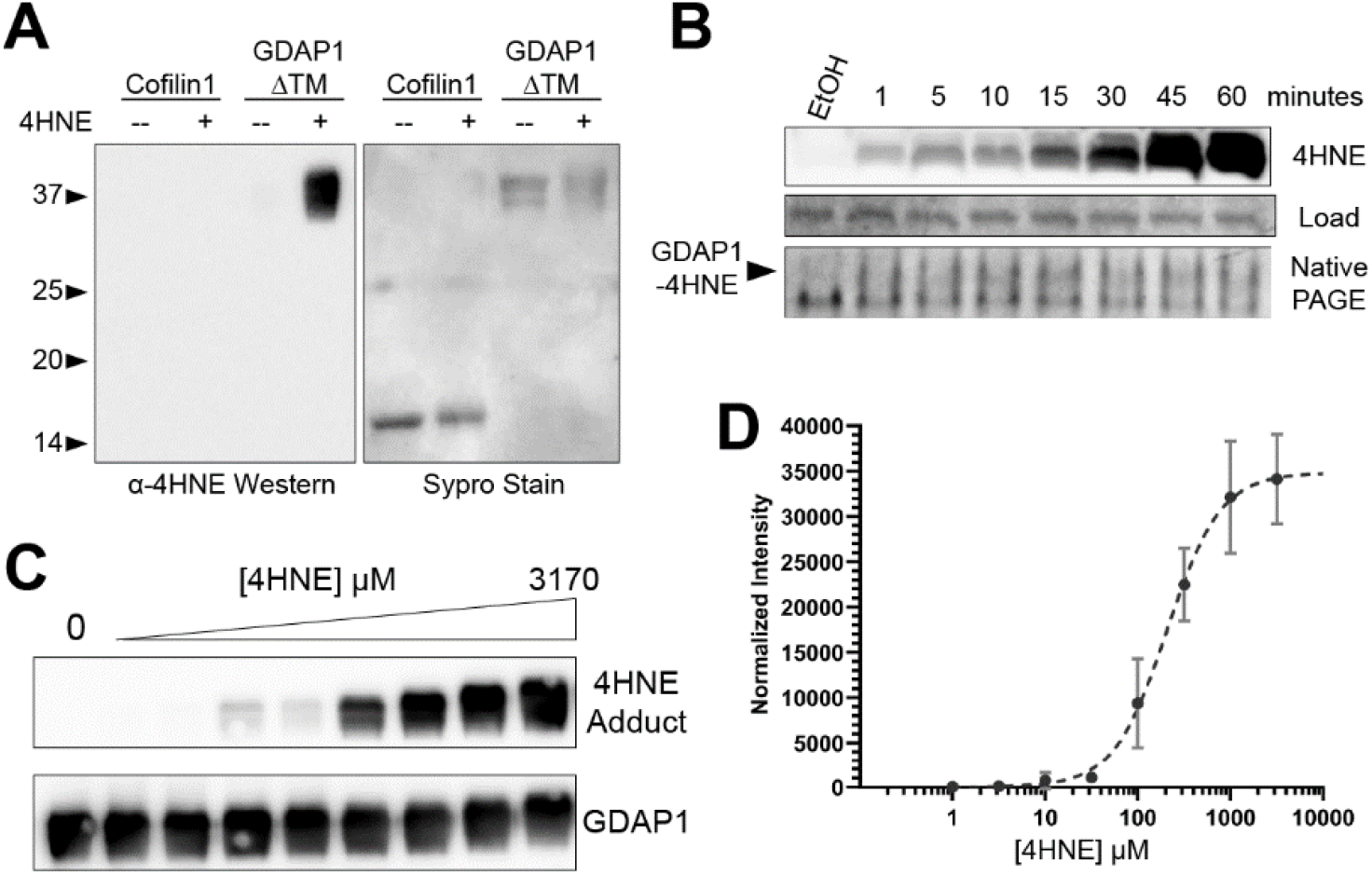
4HNE can both bind and modify GDAP1. **A)** Formation of 4HNE adducts on GDAP1 as detected by α-4HNE Western, **B)** time course of interactions between 4HNE and GDAP1. Adduct formation is monitored via anti-4HNE Western, binding by native PAGE and Sypro staining as a loading control for the Western **C)** Concentration dependence of 4HNE adduct formation. **D)** Quantification of C. Error bars represent standard deviation from three independent experiments.

### Binding to 4HNE requires the α-Loop domain in GDAP1

We next asked which domains within GDAP1 are needed to support 4HNE binding and/or adduct formation. We tested adduct formation using GDAP1ΔTM as our point of reference, and then tested further deletion of the N-terminal extension, HD1, both HD1 and NT deletions, and deletion of the α-loop. A diagram of the domain boundaries and organization is shown in **Figure 4A**. Our results demonstrate that deletion of the α-loop results in a severe reduction in 4HNE adduct formation (**Figure 4B**). This suggests that either the modification site is found within the α-loop and/or the α-loop contains residues critical for 4HNE binding. Deletion of the α-loop also resulted in a loss of 4HNE binding as detected by native PAGE (**Figure 4C**). 4HNE binding was still observed for GDAP1 constructs lacking the N-terminal extension (NT) and HD1 domains, suggesting those domains do not impact 4HNE binding (**Figure 4C**). Together these results suggest that critical determinants for 4HNE recognition reside within the α-loop.

**Figure 4.**
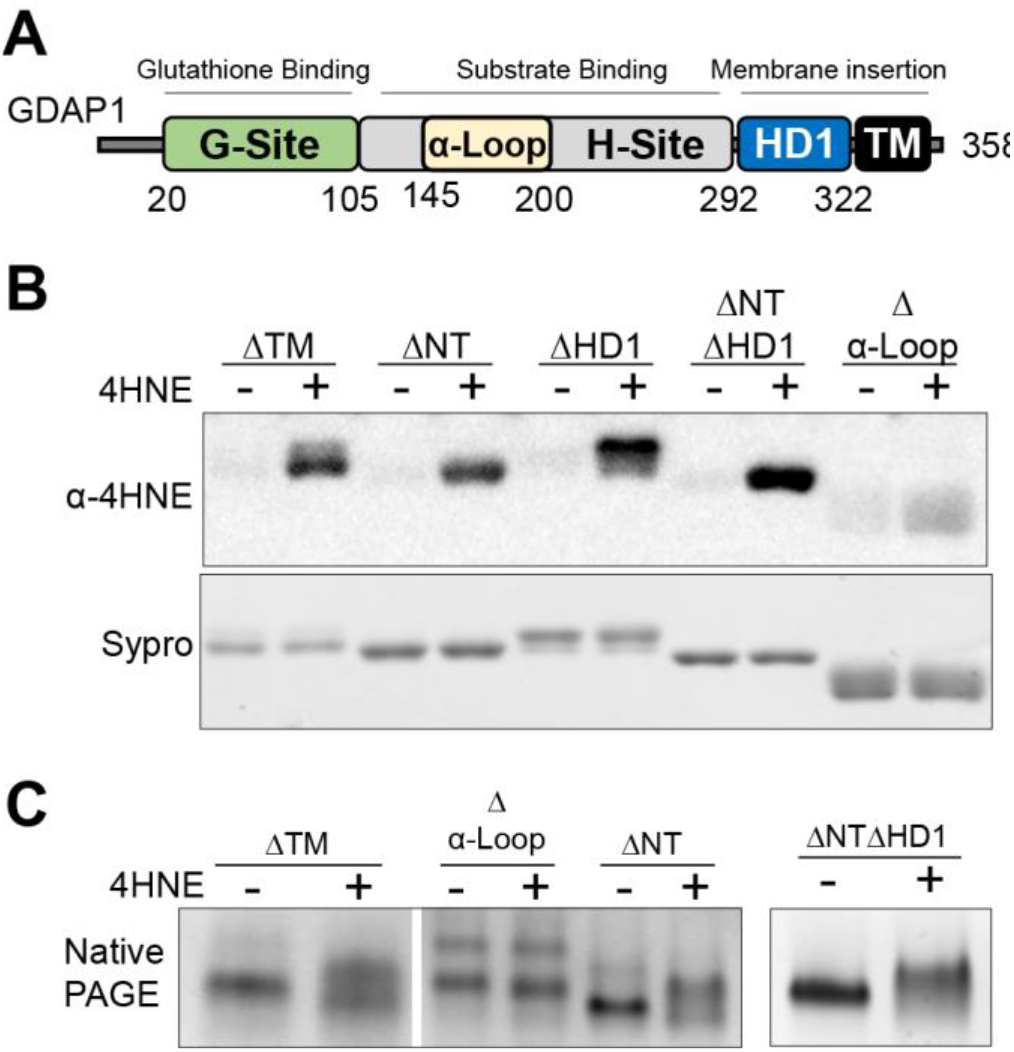
The α-loop region of GDAP1 is required for 4HNE interactions. **A)** domain organization of GDAP1 with locations of the N-terminal extension (NT, amino acids 1-20), G-site (21-108), H-site (109-282), Hydrophobic Domain 1 (HD1,292-321) transmembrane (TM, 322-358) and α-loop domain (amino acids 145-200) indicated. **B)** Adduction formation requires the α-loop. 4HNE adduct formation assay using the indicated fragment of GDAP1 probed by anti-4HNE Western as well as SYPRO staining. **C)** 4HNE binding requires the α-loop. Indicated fragments of GDAP1 incubated with or without 1mM 4HNE and analyzed by native PAGE.

### Binding of 4HNE and thapsic acid to GDAP1 are not competitive with each other

Thapsic acid has been shown to bind in an H-site pocket distinct from the canonical GST binding pocket with a binding affinity of 45 μM (Nguyen et al., 2020) (**Figure 6A**). As the GDAP1 H-site contains both 4HNE and thapsic acid binding capacity, we asked whether these molecules compete for the same binding site. Since thapsic acid interacts with GDAP1 with higher affinity than 4HNE, we anticipated that thapsic acid would effectively compete with 4HNE for adduct formation if the two ligands are utilizing the same binding sight. We tested adduct formation under these conditions and found no significant change in levels of 4HNE adduct formation in the presence of thapsic acid, suggesting that access to the site of adduct formation was not altered by the addition of thapsic to the reaction (**Figure 5A**). Next, we addressed the impact of 4HNE and thapsic acid binding on the GDAP1 protein. To do this, we N-terminally labeled GDAP1 with fluorescein and monitored the effect of adding 4HNE or thapsic acid into the reaction using fluorescence polarization. We first measured fluorescence polarization values throughout a titration of 4HNE (**Figure 5B**), finding an increase in polarization that is concentration dependent, consistent with an increase in the radius of gyration for GDAP1 upon 4HNE addition. Deletion of the α-loop largely blocks this effect, consistent with our earlier findings that the α-loop is required for 4HNE binding (**Figure 5B**). A similar titration with thapsic acid demonstrated a smaller but reproducible decrease in polarization values, suggesting that the protein’s radius of gyration is decreasing upon thapsic acid binding. The K_D_ for this effect is ~34μM, which is consistent with previous measurements of thapsic acid recognition in its binding pocket (Nguyen et al., 2020). Performing the experiment with GDAP1Δα-loop resulted in nearly identical binding curves, demonstrating that the α-loop has no impact on thapsic acid binding in this experimental context (**Figure 5C**). Next, we pre-incubated GDAP1ΔTM with 150 μM thapsic acid and asked whether this could compete with 4HNE for binding to GDAP1. As compared to the vehicle control, thapsic acid pre-incubation resulted in no apparent change in the ability of GDAP1 to bind 4HNE effect (**Figure 5D**). Pre-incubation of GDAP1ΔTM with 1 mM 4HNE for 15 minutes (conditions in which adduct formation was minimal), followed by a thapsic acid titration showed that the addition of thapsic acid could not diminish the 4HNE mediated increase in polarization (**Figure 5E**). From this data, we conclude that the 4HNE and thapsic binding sites are separate, have opposing impacts on GDAP1 structure, and that binding of thapsic acid has no impact on 4HNE binding.

**Figure 5.**
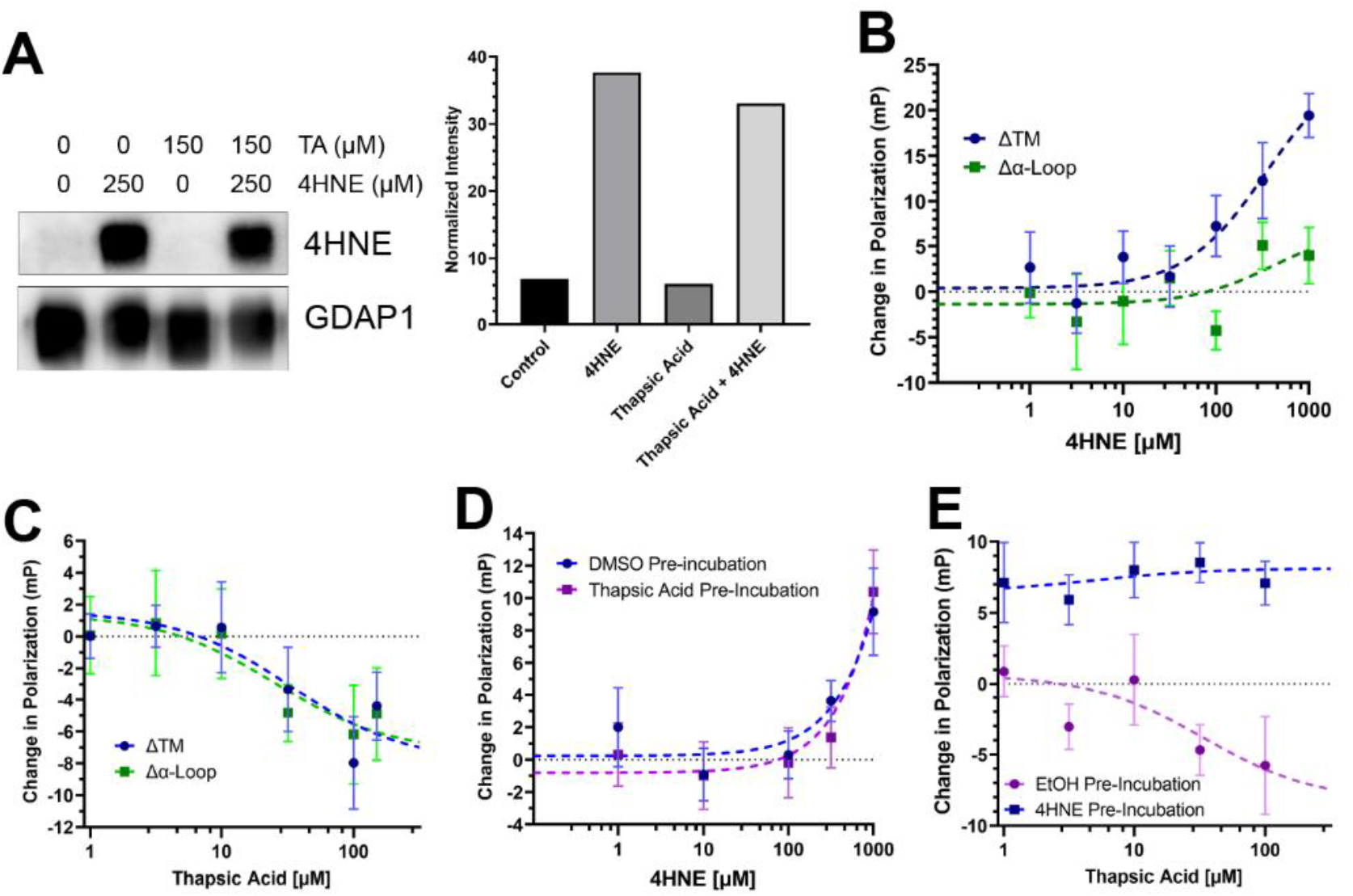
Binding of 4HNE and Thapsic acid to GDAP1 are independent. **A)** 4HNE adduct formation on GDAP1, as monitored by 4HNE Western, is not affected by pre-incubation with thapsic acid. **B)** Fluorescence polarization as a function of 4HNE concentration. An α-Loop deletion construct shows a loss in 4HNE dependent signal. P<0.0001 **C)** An α-Loop deletion shows no change in its ability to interact with thapsic acid, P=0.96. **D)** Pre-incubation of GDAP1ΔTM followed by 4HNE titration, shows that the presence of thapsic acid does not change the affinity of GDAP1 for 4HNE, P=0.38. **E)** Thapsic acid could not outcompete the effect of 4HNE binding, P<0.0001.

### Structure of GDAP1ΔTM shows the CMT mutant T157P alters α-loop positioning

Over a hundred separate missense mutations in *GDAP1* have been identified in patients with the GDAP1-type Charcot-Marie-Tooth (CMT) disease (Ammar et al., 2003; Marco et al., 2004; Martin et al., 2015; Nelis et al., 2002; Rzepnikowska and Kochanski, 2018; Senderek et al., 2003). Both dominant and recessive mutations have been identified and mutations have been classified into CMT sub-types based on their effect on myelination and nerve conduction velocities (Senderek et al., 2003). With regard to the structure of GDAP1, many CMT mutations have been identified that strongly cluster on long helices 4 and 5 in a region called the CMT hotspot (Googins et al., 2020). Recent structural data on GDAP1 containing two CMT-causing mutants in the CMT hotspot demonstrate that neither alters protein stoichiometry, but instead disturb a network of intermolecular interactions that alter protein stability (Sutinen et al., 2022). Mutations outside of this CMT hotspot would not be expected to impact this interaction network and have not been explored biochemically or structurally. Towards this end, we crystallized and determined the structure of GDAP1ΔTM T157P. Details of the structure determination process are described in detail in the Experimental Methods. The crystals contain a pair of GDAP1 proteins in the asymmetric unit in a disulfide linked arrangement similar to that observed for CMT-causing mutants H123R and R120W (Sutinen et al., 2022). Both proteins are ordered throughout the GST-core which aligns with an r.m.s.d of 0.72 Å over 171 Cα atoms. Electron density in loop regions was generally better for chain B which was used for the analysis and figures presented here. Differences between GDAP1-T157P and other GDAP1 structures (experimental and predicted) are focused within two regions: the α-loop and helix α2 of the G-site. As defined in the literature the α-loop is a sequence motif containing GDAP1 residues 145-200 (Estela et al., 2011; Shield et al., 2006) that is inserted between helices α4 and α5 in the H-site (**Figure 6A**). Within the α-loop, residues 154-200 are predicted to fold into a two-helical element (described as helices α4a α5a here), flanked by hinge regions (described here as N- and C-hinges) (**Figure 6B**) believed to promote transition of the GDAP1 α-loop between closed and open states(Sutinen et al., 2022). The CMT-causing T157P mutant is located very close to the N-hinge (approximately residue 154), where the rigidity of the introduced proline might be expected to alter α-loop motions (**Figure 6B**). Alternatively, since helix α4a has already been demonstrated to be highly flexible (Googins et al., 2020; Nguyen et al., 2020; Sutinen et al., 2022), the proline substitution may simply add destabilize helix α4a further, leaving helix α5a unaltered. Here we find that both helices α4a and α5a are disordered, suggesting that changes in the N-hinge affect positioning of the entire α-loop. This is consistent with a concerted motion of helices α4a and α5a between an extended or “open” position observed previously and a “closed” position as predicted by AlphaFold2.

**Figure 6.**
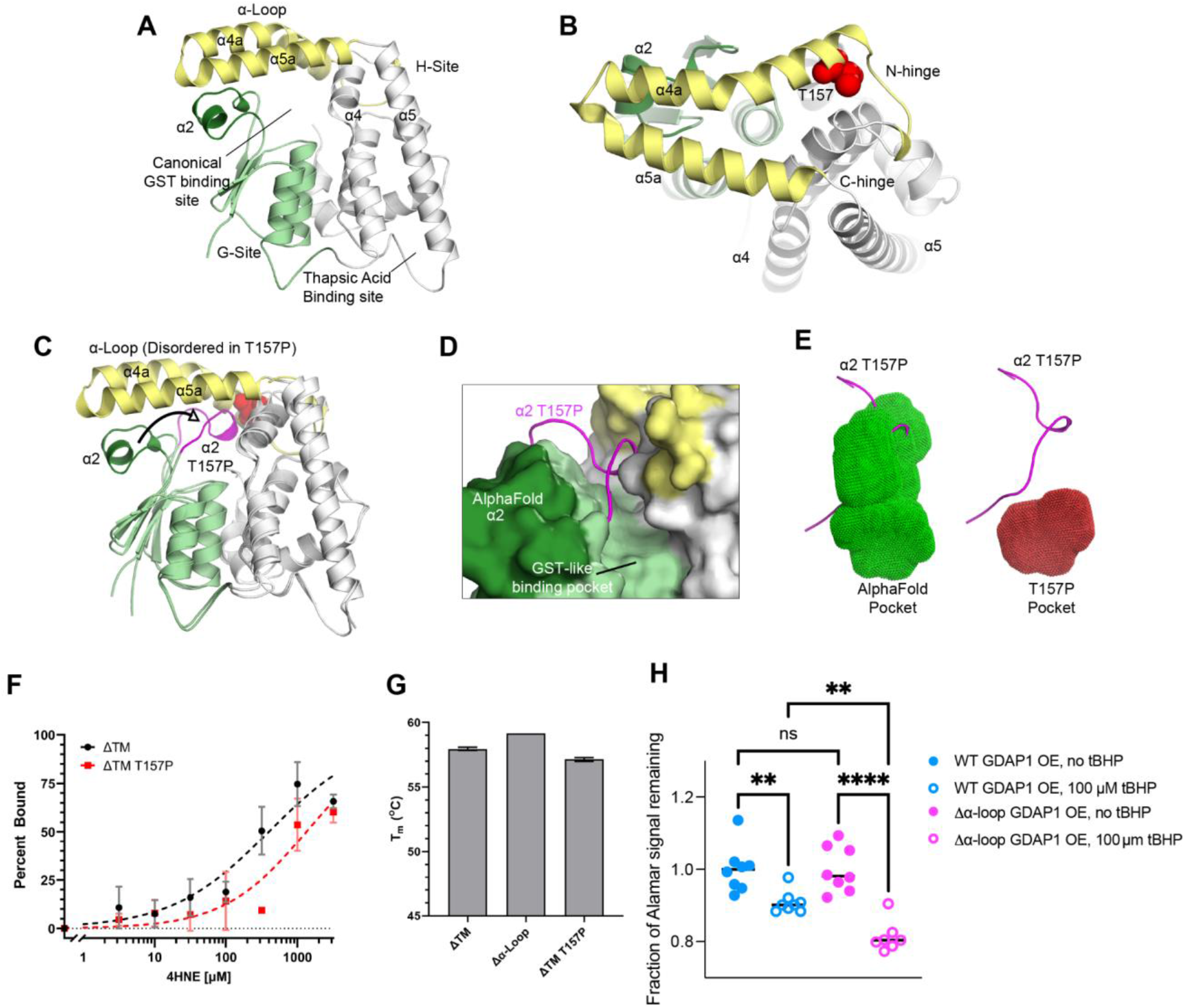
CMT-causing mutant T157P destabilizes the α-loop, facilitating motions in α2, decreasing 4HNE binding affinity and GDAP1 cytoprotective function. **A)** Predicted structure of GDAP1 generated from AlphaFold2, colored as in Figure 4A with HD1 and TM domains omitted for clarity. **B)** Top view of A, highlighting the position of T157 (Red spheres) and its proximity to the N-hinge region. **C)** Structural overlap of AlphaFold and experimental T157P structures. Both colored as in 4A, except for G-site helix α2 which is dark green in the predicted structure and magenta in the T157P structure. **D)** α2 occupies most of the canonical GST binding pocket in the T157P structure. Surface shown is from the AlphaFold prediction with most of the alpha-loop removed for clarity. **E)** Comparison of the canonical binding pocket between WT (green) and T157P (red) structures. Pocket volume visualized as dots generated by 3V analysis. The position of the α2 helices from the structural overlap of the T157P structure is shown in magenta. **F)** 4HNE binding isotherms for GDAP1ΔTM and GDAP1ΔTM-T157P, demonstrate a reduction in 4HNE binding affinity. **G)** Melting temperature of GDAP1ΔTM, Δα-Loop, and T157P constructs are relatively similar as assessed by differential scanning fluorometry. **H)** Comparting the effects of tBHP on AlamarBlue fluorescence in cells overexpressing WT (GDAP1 OE) and △α-loop GDAP1. Compared to WT GDAP1, Cells that overexpress △α-loop GDAP1 offer diminished protection against tBHP, which is indicated by fact that following tBHP application, a significancy smaller fraction of Alamar signal remains in cells overexpressing △α-loop GDAP1 relative to WT GDAP1. Since Alamar signal is lost in oxidizing environments, this indicates a more oxidizing environment in tBHP-treated △α-loop GDAP1 overexpressing cells relative to WT GDAP1 overexpressing cells **: p<0.001, ***: p<0.0001.

**Table 1.**
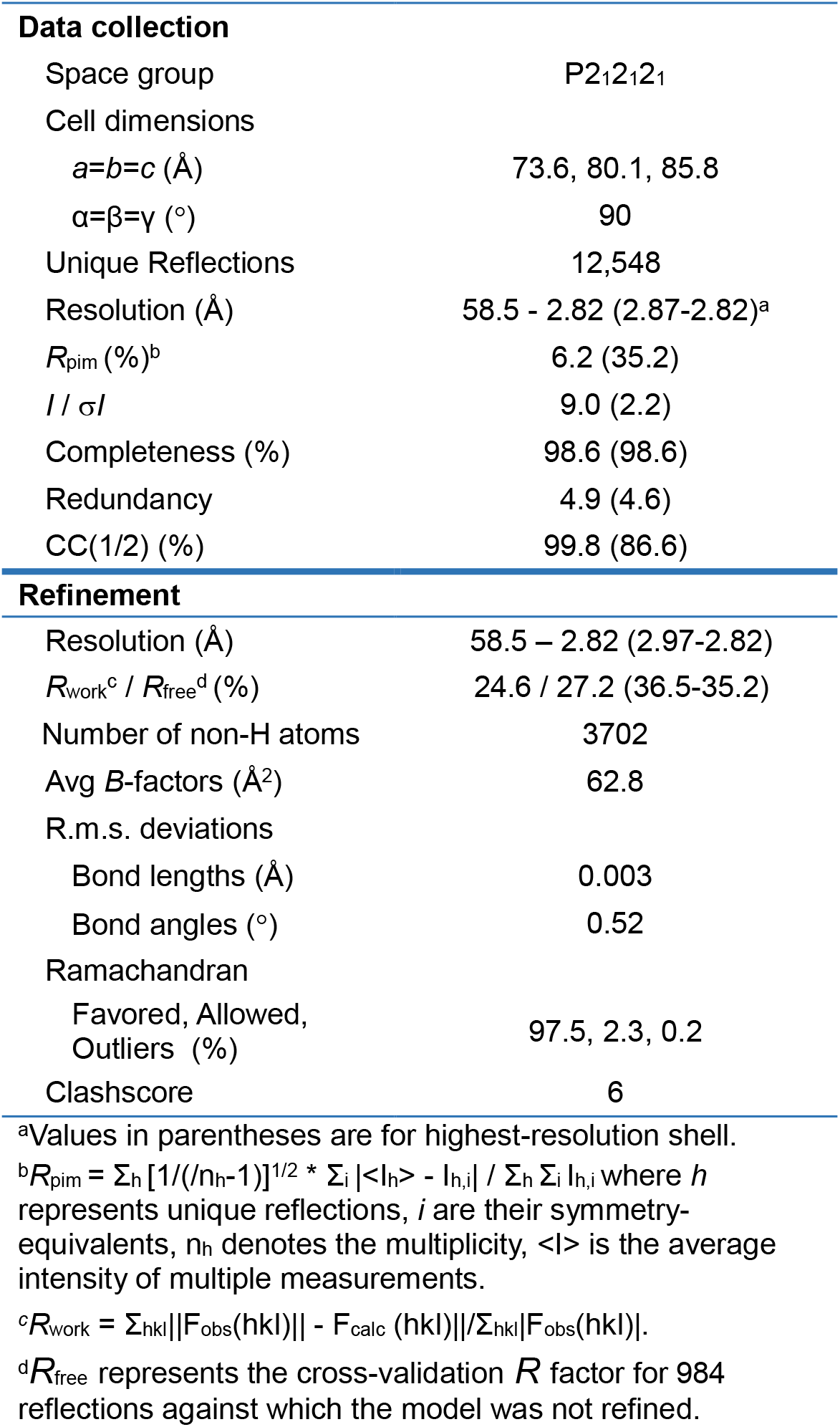
Data Collection and Refinement.

|  |  |
| --- | --- |
| Space group | P2<sub>1</sub>2<sub>1</sub>2<sub>1</sub> |
| Cell dimensions |  |
| $a=b=c$ (Å) | 73.6, 80.1, 85.8 |
| $\alpha=\beta=\gamma$ (°) | 90 |
| Unique Reflections | 12,548 |
| Resolution (Å) | 58.5 - 2.82 (2.87-2.82)<sup>a</sup> |
| $R_{\text{pim}}$ (%)<sup>b</sup> | 6.2 (35.2) |
| $I / \sigma I$ | 9.0 (2.2) |
| Completeness (%) | 98.6 (98.6) |
| Redundancy | 4.9 (4.6) |
| CC(1/2) (%) | 99.8 (86.6) |
| **Refinement** |  |
| Resolution (Å) | 58.5 – 2.82 (2.97-2.82) |
| $R_{\text{work}}^c / R_{\text{free}}^d$ (%) | 24.6 / 27.2 (36.5-35.2) |
| Number of non-H atoms | 3702 |
| Avg $B$-factors (Å<sup>2</sup>) | 62.8 |
| R.m.s. deviations |  |
| Bond lengths (Å) | 0.003 |
| Bond angles (°) | 0.52 |
| Ramachandran |  |
| Favored, Allowed, Outliers (%) | 97.5, 2.3, 0.2 |
| Clashscore | 6 |

<sup>a</sup>Values in parentheses are for highest-resolution shell.
<sup>b</sup> $R_{\text{pim}} = \sum_h [1/(n_h-1)]^{1/2} * \sum_i |<I_h> - I_{h,i}| / \sum_h \sum_i I_{h,i}$ where $h$ represents unique reflections, $i$ are their symmetry-equivalents, $n_h$ denotes the multiplicity, $<I>$ is the average intensity of multiple measurements.
<sup>c</sup> $R_{\text{work}} = \sum_{hkl} ||F_{\text{obs}}(hkl)| - F_{\text{calc}}(hkl)| / \sum_{hkl} |F_{\text{obs}}(hkl)|$ .
<sup>d</sup> $R_{\text{free}}$ represents the cross-validation $R$ factor for 984 reflections against which the model was not refined.

In canonical GST enzymes, helix α2 in the G-site forms one wall of the active site and functions in glutathione recognition. In GST P1, this element has been shown to be flexible and to undergo motions concomitant with binding of its substrates (Lo Bello et al., 1998; Oakley et al., 1998). Structural predictions from AlphaFold position helix α2 in an orientation similar to the bound state of GST P1. In GDAP1 T157P, however, we find helix α2 in an extended position, deep within the canonical binding pocket (**Figure 6C and 6D**). This results in an ~70% decrease in the volume of the binding pocket using the 3V cavity calculator (Voss and Gerstein, 2010) (**Figure 6E**). We hypothesized that this dramatic change in the pocket would result in a decrease in binding affinity between 4HNE and GDAP1 T157P. We tested this directly by purifying recombinant GDAP1ΔTM T157P protein and assessing its ability to bind 4HNE via native PAGE under conditions with minimal adduct formation (**Figure 6F**). The binding affinity for this interaction is measured at 1330 ± 259 μM. From this we can conclude that either, 1) 4HNE interacting residues are housed within the canonical GST binding pocket that these interactions are lost by movement of helix α2 into the pocket, or 2) 4HNE interacting residues are contained within the α-loop and the failure to adopt the “closed” conformation of GDAP1 results in a decrease in binding. The degree to which one or both of these conclusions is contributing to 4HNE binding is currently unknown. Lastly, we performed protein thermal shift assays on the GDAP1ΔTM T157P mutant and found that its thermal transition was very similar to that of wild-type, demonstrating that the loss of 4HNE binding is not the result of an overall loss of stability or folding (**Figure 6G**). These data all suggest that the T157P variant of GDAP1 has a defect in 4HNE recognition. To validate the role of α-loop in the protective function of GDAP1, we used HEK293 cells stably overexpressing a GFP-tagged GDAP1 α-loop deletion construct (**Figure 6H**). Expression was confirmed using GFP fluorescence, and the ability of cells to resist redox changes induced by the prooxidant tBHP was analyzed using AlamarBlue as discussed above. Compared with wild-type GDAP1, cells that overexpress a GDAP1 α-loop deletion construct show significantly reduced protection against tBHP. Together, both our in vitro and cell-based assays show that the α-loop region of GDAP1 plays a critical role in 4HNE recognition and GDAP1’s cytoprotective function in the cell.

## Discussion

GDAP1 activity has been implicated in a variety of mitochondrial functions, including the redox homeostasis (Noack et al., 2012), the oxidative stress response (Niemann et al., 2014), and the regulation of mitochondrial network dynamics (Googins et al., 2020; Huber et al., 2016; Niemann et al., 2005; Pedrola et al., 2008; Pedrola et al., 2005). While the involvement within these pathways seems apparent, a direct molecular connection that establishes a role for GDAP1 has remained elusive. Structural data and sequence homology have all suggested that GDAP1 contains significant similarity to the GST family of detoxifying enzymes (Marco et al., 2004; Shield et al., 2006), however it adopts a non-canonical quaternary structure (Nguyen et al., 2020), contains unique GDAP1-specific domains, and a role for its active site has not been established (Googins et al., 2020). Here, we have presented biochemical, structural, and cell-based analyses to define the role of the canonical GST-like binding pocket in GDAP1 function.

We focused on GDAP1’s role as a member of the oxidative stress response pathway as this pathway is well established for the GST superfamily of which GDAP1 is a novel member (Marco et al., 2004; Shield et al., 2006). In this context, we asked whether GDAP1 could interact with the end products of lipid peroxidation, specifically MDA and 4HNE which are both toxic end-products formed by the decomposition of oxidized and peroxidated lipids under conditions of oxidative stress (Esterbauer et al., 1991; Poli et al., 2008). We determined that GDAP1 specifically recognized 4HNE and that this binding occurs within the canonical GST- like binding pocket. We revealed that the α-loop, a sequence insertion within the H-site that is specific to GDAP1, was required for 4HNE binding. Using xray crystallography, we reveal that disease-causing mutants within the N-hinge region destabilize positioning of GDAP1’s α-loop with a concomitant movement of the α2 helix from the G-site into the binding pocket, and establish a connection between these two critical regions of the protein and 4HNE binding. GDAP1 has also been shown to interact with the lipid thapsic acid (Nguyen et al., 2020), however, we demonstrate here that these binding events occur at separate locations and do not appear to be allosterically linked to each other. The biological role for thapsic acid in GDAP1 function is intriguing but remains unclear.

The establishment of 4HNE as a GDAP1 binding partner is a significant breakthrough in GDAP1 biology that unlocks several intriguing mechanistic questions. First, we observe both covalent and non-covalent interactions with 4HNE. Our results suggest a role for GDAP1 in responding to 4HNE levels but it is unclear if both modes of interaction participate biologically. It is possible that the slower-forming but longer lasting adduct formation provides a timing mechanism following conditions of oxidative stress. Alternatively, it could modulate GDAP1 activity, either positively or negatively, until the 4HNE-adduct can be reversed by increased glutathione levels when redox homeostasis is restored. The mechanism of 4HNE binding and its molecular and cellular consequences are open questions that we are actively pursuing.

Many CMT-causing mutations have been identified within the “CMT hotspot” located on long helices 4 and 5 (Cassereau et al., 2011a; Cassereau et al., 2011b; Googins et al., 2020; Rzepnikowska and Kochanski, 2018). Mutants in the hotspot have been suggested to influence the thapsic acid binding region or to alter overall stability (Nguyen et al., 2020; Sutinen et al., 2022). Here we present biochemical and structural data on T157P a mutant within the N-hinge region, which is distant and distinct from the CMT hotspot. Our data indicate that changes in positioning of the α-loop and α2 regions are associated with this mutation which is reflected in changes in the 4HNE binding pocket and GDAP1 binding affinity for 4HNE. This data provides a molecular explanation for the capacity of T157P to promote disease and we hypothesize that other CMT causing mutants in the binding pocket (such as S34, S36 or K39) or within the α-loop itself (P153, K161, N178, or L205) might have a similar 4HNE binding defects. Future work will explore the direct connection of these residues to interactions with 4HNE.

## Materials and Methods

### Protein Purification

Coding sequences for mouse *GDAP1* constructs encoding GDAP1ΔTM, GDAP1ΔTM T157P, GDAP1ΔαL, and all other GDAP1 constructs described here, were PCR amplified and cloned into the pKF3 plasmid (Googins et al., 2020) for bacterial expression with an N-terminal His10-mRuby2 tag which can be removed by cleavage with TEV protease. The resulting proteins retain GGS on their N-terminus. Expression was performed in BL21(DE3)- RIPL *Escherichia coli* cells (Agilent) in LB at room temperature and induced through the addition of 0.2 mM IPTG for ~24 hours. Cells were harvested, resuspended in [200 mM NaCl, 20 mM Tris pH8, 40 mM Imidazole pH8, 1 mM Tris(2-carboxyethyl) phosphine (TCEP), 5% glycerol], and lysed by homogenization (Avestin C-3). Insoluble material was removed by centrifugation at 16,000 x *g* and His_10_-mRuby2-GDAP1 fusion protein captured using nickel affinity chromatography followed by digestion with TEV protease overnight to liberate GDAP1 protein from the His10-Ruby tag. A second round of nickel affinity chromatography was then performed to remove the mRuby2-tag and TEV. Additional purification was achieved through anion exchange chromatography, and a final step of gel filtration was used to remove any potential aggregates and lingering contaminants. ThermoFluor of samples indicate a single-phase thermal transition consistent with a folded protein. Coding sequences for Cofilin1 were also cloned into pKF3 and expressed and purified in manner similar to GDAP1.

### Native PAGE protein-ligand interaction assays

Analysis of protein-ligand interactions via Poly-acrylamide Gel Electrophoresis was conducted by mixing potential ligands (Indicated ligand at indicated concentration) with GDAP1 protein at a final concentration of 55 μM. The reaction buffer was 50mM Tris pH 8, 50mM NaCl. Incubation times were 15 minutes at room temperature unless indicated. Reactions were loaded using 4x Native loading buffer [200 mM Tris pH 6.8, 50% glycerol] supplemented with 0.5mM β-mercaptoethanol to minimize non-specific 4HNE-protein adducts and run on 12% PAGE for 3.5 hours at 200 volts. Gels were stained with Coomassie stain and imaged using an Amersham Imager 600 (General Electric) transillumination setting and analyzed using FIJI for band intensity. Gels imaged using Sypro Orange were incubated in 7.5% acetic acid with 0.05% SDS for 30 minutes, washed with 7.5% acetic acid, then stained with 2x SYPRO Orange dye (Invitrogen) in 7.5% acetic acid for 30 minutes. Sypro stained gels were imaged using an Amersham Imager 600 (General Electric) and analyzed with FIJI-ImageJ (Schindelin et al., 2012).

### 4HNE modification reactions

Measurements of 4HNE adduct formation were performed with cofilin1 or the indicated GDAP1 fragment at a final protein concentration of 5 μM in reaction buffer consisting of [50mM Tris pH 8, 50mM NaCl]. Cofilin1 and GDAP1ΔTM reaction was incubated with 1 mM 4HNE for 2 hours at room temperature in a manner similar to previous literature regarding cytochrome C modification (Isom et al., 2004). The exception is during the time course experiment in which reaction times are indicated. Samples were mixed 3:1 with 4x loading buffer (200 mM Tris pH 6.8, 8% SDS, 50 mM EDTA, 0.8% Bromophenol Blue, 50% glycerol) without reducing reagent to preserve 4HNE adducts, and boiled at 100 °C for 5 minutes. 200 ng of protein loaded in each well and run on a 15% SDS-PAGE for Western Blot analysis. The SDS-PAGE gel was transferred onto nitrocellulose membrane (Thermo Scientific) that was then treated with 100 mM sodium borohydride (Fisher Chemical) for 20 minutes to stabilize 4HNE adducts (McCormack et al., 2005). Western blotting was performed using a α-4HNE antibody specific to the Michael protein adduct (EMD Millipore Corp.) at a dilution of 1:1000 antibody over night at 4°C. Goat α-Rabbit HRP (Thermo Scientific) was the secondary antibody (1:3000 dilution) for 1 hour at room temperature with intervening wash steps performed with TBST. Modified protein was detected by using SuperSignal West Pico PLUS Chemiluminescent Substrate kit (ThermoScientific) with an Amersham Imager 600 (General Electric). Blots were then stripped and re-probed with goat α-GDAP1 (Sigma) polyclonal antibody (1:1000 dilution) overnight at 4°C, followed by rabbit anti-goat HRP secondary antibody (1:3000) for 1 hour at room temperature, with intervening TBST washes and imaged as above. Band Intensities were quantified using FIJI-ImageJ. 4HNE band intensity was normalized against the GDAP1 band intensities. Results were analyzed using PRISM.

### Fluorescein Labeling of Proteins

The indicated GDAP1 constructs were N-terminally labeled with fluorescein as in (Mohan et al., 2013). Briefly, protein was dialyzed into fluorescein labeling buffer (20 mM HEPES pH 7.0, 100 mM NaCl, 8% glycerol) at a final protein concentration of 140 μM, then incubated with 10x molar excess of Fluorescein (Invitrogen) for 2 hours at room temperature. Protein was then dialyzed against (20 mM HEPES pH 8, 200 mM NaCl, 2% Glycerol, 1 mM TCEP) for 4 days with a buffer changes every day, until no free fluorescein could be detected by either PAGE or spectroscopically.

### Fluorescence Anisotropy

Analysis of GDAP1 via fluorescence anisotropy was conducted using fluorescein labeled protein at 50 nM supplemented with unlabeled protein to achieve a final concentration of 12.5 μM. The reaction buffer was 50 mM Tris pH 8 and 50 mM NaCl. Unless indicated, protein was incubated for 30 minutes with the indicated ligand prior to measurement of fluorescence polarization using a Biotek Cytation 5 imaging reader.

### Crystallography

GDAP1ΔTM T157P was purified as described earlier and stored at −80°C prior to crystallization trials. Crystals were obtained using sitting-drop vapor diffusion method at 4°C. 1 μL of protein at 12.8 mg/ml was added to 1 μL of well solution containing 0.2 M Ammonium Sulfate, 0.1 M Bis-Tris pH 5.5, and 25% (w/v) PEG3350. Small amorphous looking crystals grew slowly over the course of 6 months. Crystals were soaked in mother liquor supplemented with 15% glycerol and flash frozen in liquid nitrogen prior to data collection.

Diffraction data were collected at beamline 31-IDD at Argonne National Labs and processed and scaled to 2.8 Å resolution via AutoPROC (Vonrhein et al., 2011) using *I/σI* >2.0 and CC(1/2) >0.3 as cutoffs. Crystals of GDAP1ΔTM T157P belong to space group P2_1_2_1_2_1_ with *a*=73.59, b=80.06, and c=85.81 Å. Phases were estimated via the molecular replacement method using the structure of the GST-like core of GDAP1 as the search model (Googins et al., 2020) (PDBID:6UIH). An initial model was built into density using COOT(Emsley et al., 2010) and further improved through rounds of refinement in Phenix(Adams et al., 2010) and model building in COOT, including simulated annealing in the first refinement step. Positional and group B-factor refinements were used during this process. Model quality was assessed using MolProbity within Phenix. Model and structure factors files for the GDAP1 -T157P are deposited in the PDB under PDBID code 8EXZ.

### Cell culture and stable lines

HEK293 cells were grown in a 5% CO_2_ humidified atmosphere at 37°C in Dulbecco-modified eagle medium supplemented with 10% fetal bovine serum. To generate stable lines, the cells were transfected with cDNA plasmids using the calciumphosphate method and, 24 hours post-transfection, seeded at low density into 400 mg/ml G418. Colonies were picked by scraping and subcloned into 24-well plates. The expression and transfection rate were confirmed using fluorescent microscopy. Alternatively, the cells were transiently transfected using the calcium-phosphate method and used for experiments within 24 to 48 hours post-transfection.

### Fluorescent measurements in live cells

The cells were seeded into 96-well plates at high confluency and DMEM was replaced with HEPES-based buffer containing, in mM: 140 NaCl, 5 KCl, 1 MgCl_2_, 1 CaCl_2_, 10 HEPES at pH 7.4 and supplemented with 1 g/l glucose. The AlamarBlue (ThermoFisher Scientific, Waltham, MA, product number DAL1025) staining was performed according to the manufacturer’s instructions: 10 μl of the reagent was added per well and the reading of 560/590 nm fluorescence commenced, using ThermoFisher Fluoroscan plate reader. The data were read every 15 min for 3 hours and each well’s readings were normalized to the value recorded at the beginning of the read. The conversion of AlamarBlue into a fluorescent product is suppressed if the environment of the cytoplasm is oxidizing; to quantitatively compare the effects of GDAP1 overexpression and drug application of cytoplasmic redox, all data points were normalized to corresponding time points and concentrations in control, untreated cells.

WST-1 measurements were performed per manufacturer’s (Sigma Aldrich, St Louis, MO, product number 11644807001) instructions: 10 μl of the reagent was added per well, and the 340 nm absorbance was read using Accuris SmartReader 96 (Accuris Instruments, Edison, NJ). To induce a shift in the cellular redox, the cells were treated with 50-500 μM tert-Butyl hydroperoxide (tBHP) for 1 hr before the experiment. The data are presented as the percentage of AlamarBlue signal that is lost as a result of tBHP application.

### Statistical analysis

Ligand binding and modification curves were fitted using the Specific Binding with Hill slope function of Prism 9 and response curves to ligand interactions obtained from fluorescence polarization assays were fitted to data using the Dose Response (3 Parameters) function of Prism 9. It is from these curves that K_D_ and EC_50_ values were calculated with corresponding SEM variance values. Comparisons of these curves were done using sum-of-squares F test with null hypothesis that all data can be fitted using one curve. If the null hypothesis was rejected at p<0.05, we concluded that the curves were statistically different from each other. AlamarBlue concentration curves were fitted using the Michaelis-Menten function of Prism 9 and compared using extra sum-of-squares F test with null hypothesis that all data can be fitted using one curve. If the hypothesis was rejected at p<0.05, we concluded that the curves are significantly different from each other. Alternatively, pairwise comparison (AlamarBlue and WST-1) was performed using ordinary one-way Anova with multiple comparisons by means of Bartlett’s test. P<0.05 was considered significant.

## Acknowledgements

This research used resources of the Advanced Photon Source, a U.S. Department of Energy (DOE) Office of Science User Facility operated for the DOE Office of Science by Argonne National Laboratory under Contract No. DE-AC02-06CH11357. Use of the Lilly Research Laboratories Collaborative Access Team (LRL-CAT) beamline at Sector 31 of the Advanced Photon Source was provided by Eli Lilly Company, which operates the facility. This work was supported by NIH R21 grant NS094860 (K.K.).

## Author Contributions

A.P.V. and K.K. designed experiments, analyzed the data, and wrote the paper. M.R.G. performed the crystallization and structure determination, as well as the biochemical experiments. A.P.V. analyzed the results of the structural and biochemical experiments. M.R.B., A.O.W. and K.K. performed the cell-based experiments. K.K. analyzed the results of the cell-based experiments.

